# The B-type cyclin Clb4 prevents meiosis I sister centromere separation in budding yeast

**DOI:** 10.1101/2024.12.18.629243

**Authors:** Gal Lumbroso, Gisela Cairo, Soni Lacefield, Andrew W. Murray

**Affiliations:** Department of Molecular and Cellular Biology, Harvard University, Cambridge MA, USA; Department of Biochemistry and Cell Biology, the Geisel School of Medicine at Dartmouth, Hanover NH, USA

## Abstract

In meiosis, one round of DNA replication followed by two rounds of chromosome segregation halves the ploidy of the original cell. Accurate chromosome segregation in meiosis I depends on recombination between homologous chromosomes. Sister centromeres attach to the same spindle pole in this division and only segregate in meiosis II. We used budding yeast to select for mutations that produced viable spores in the absence of recombination. The most frequent mutations inactivated *CLB4*, which encodes one of four B-type cyclins. In two wild yeast isolates, Y55 and SK1, but not the W303 laboratory strain, deleting *CLB4* causes premature sister centromere separation and segregation in meiosis I and frequent termination of meiosis after a single division, demonstrating a novel role for Clb4 in meiotic chromosome dynamics and meiotic progression. This role depends on the genetic background since meiosis in W303 is largely independent of *CLB4*.

## Introduction

Meiosis is a specialized cell division in which one round of DNA replication, followed by two rounds of chromosome segregation halves a cell’s ploidy. Mitosis and the second meiotic division (meiosis II) segregate sister centromeres, but they remain attached to each other in meiosis I. Instead, meiosis I segregates homologous chromosomes (homologs, see figure 1A). Accurate homolog segregation requires recombination, which links the homologs to each other and directs them to attach to opposite poles of the meiosis I spindle [15]. In the budding yeast *Saccharomyces cerevisiae*, the combination of nitrogen starvation and non-fermentable carbon sources induce diploid cells to commit to meiosis and produce four haploid spores enveloped in a resilient ascus. In both yeast and humans, errors in meiosis I are a frequent source of aneuploidy.

**Figure 1.**
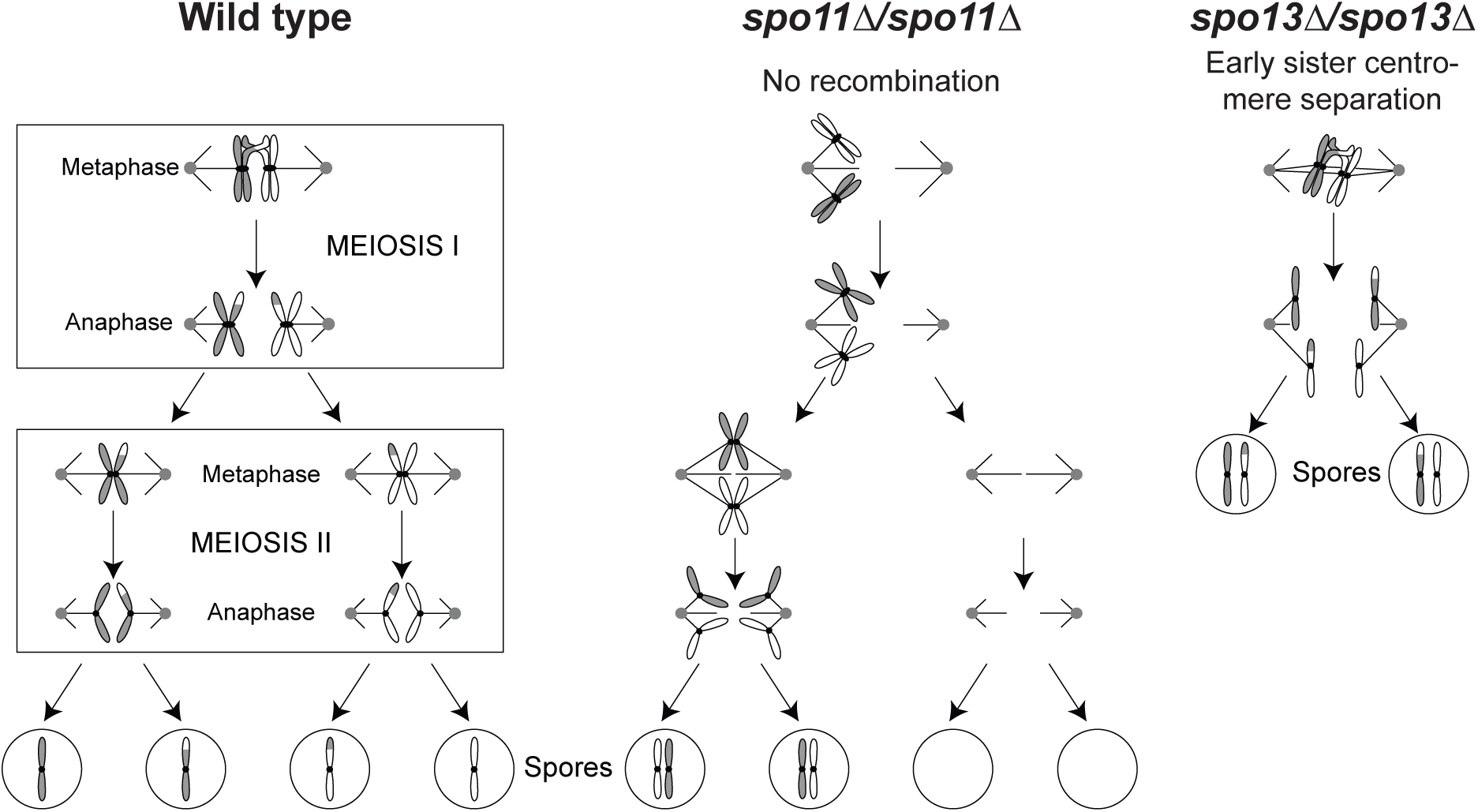
Diagram of meiosis in wild-type, mutants lacking recombination, and mutants that separate sister centromeres prematurely. **(A)** In meiosis I, homologous chromosomes (homologs) are held together by recombination. Kinetochores bound to sister centromeres are linked together (sister kinetochore mono-orientation). Homologs are segregated to opposite poles after cohesin (not shown) is cleaved at the chromosome arms. In meiosis II, kinetochores bound to sister centromeres face in opposite directions. Sister centromeres are segregated after centromeric cohesin is cleaved. **(B)** In the absence of recombination, homologs move to random poles in meiosis I, leading to aneuploidy. **(C)** Early sister centromere separation, as observed in Spo13 mutants, where sister centromeres attach to opposite poles in meiosis I. In many cases, meiosis is terminated after a single division where sisters separate, leading to the production of two diploid spores instead of four haploid spores. Accurate sister segregation does not depend on recombination, thus these mutations can rescue viability in the absence of recombination.

Meiosis I has three critical features. First, homologs are linked to each other and directed to segregate to opposite poles by recombination induced by double-strand breaks. The breaks are generated by Spo11, a conserved derivative of topoisomerase II [3, 13]. Failure to recombine, such as due to the absence of Spo11 (Fig. 1B), leads to random homolog segregation, aneuploidy, and drastically reduced gamete viability [1, 11, 15, 19, 22, 30]. Second, cohesin, the protein complex that holds sister centromeres together, is protected from cleavage near the centromere in meiosis I. This allows homolog disjunction while maintaining cohesion near the centromeres until they segregate from each other in meiosis II. Third, kinetochores, the protein complexes that assemble on the centromeric DNA and bind to microtubules, behave differently in meiosis I and II. In meiosis I, the two sister kinetochores assemble a single microtubule binding site, which attaches to only one pole of the spindle (mono-orientation), whereas in meiosis II, the two sisters attach to opposite spindle poles as they do in mitosis. The monopolin complex, which contains the four components Mam1, Csm1, Lrs4 and Hrr25 [28, 29], enforces sister kinetochore mono-orientation [25, 35]. Removing monopolin components or Spo13, a protein that regulates the activity of several protein kinases [10], allows sister centromeres to attach to opposite poles in meiosis I. In the absence of Spo13, sister centromeres segregate from each other in a single meiotic division leading to the formation of two diploid spores. (Fig. 1C).

Both meiotic divisions employ many of the same cell cycle regulators as mitosis, including Cdk1 (also known as Cdc28), which binds nine different cyclins during mitotic growth: three G1 cyclins, Cln1-3, two S-phase cyclins, Clb5 and Clb6 and four M-phase cyclins, Clb1-Clb4 [20, 21, 26, 37]. The meiotic cell cycle only employs a subset of these cyclins: the G1 cyclins are not required and Clb2 is not expressed in meiosis [6]. The apparent redundancy of M-phase cyclins in mitosis [9] gave rise to the question of whether their meiotic functions were more distinct. A study of meiosis in cyclin mutants by Dahmann and Futcher [6] reported that the triple mutant *clb1*Δ/*clb1*Δ *clb3*Δ/*clb3*Δ *clb4*Δ/*clb4*Δ drastically reduced spore formation and two double mutants (*clb1*Δ/*clb1*Δ *clb3*Δ/*clb3*Δ and *clb1*Δ/*clb1*Δ *clb4*Δ/*clb4*Δ) produced a significant fraction of asci containing only two spores (dyads), in which homologs segregated from each other as they do in a normal meiosis I. All four single cyclin deletions produced phenotypes similar to the wild-type, leaving open the question of redundancy. While Clb1 and Clb3 have since been revealed to have unique meiotic functions [4, 24], it has been unclear whether Clb4 has one as well.

We revealed a novel role of Clb4 in meiosis I chromosome dynamics by experimentally evolving *spo11*Δ/*spo11*Δ diploid cells to form viable spores in the absence of meiotic recombination. This selection produced strains that made dyads instead of tetrads and segregated sisters in their single meiotic division. The causative mutations included loss of function alleles in genes known to be required for sister cohesion in meiosis I (*SPO13* [12, 15, 36] and *MAM1* [29]) but the most frequent class were mutations that inactivated *CLB4*. The phenotype of the *CLB4* mutations did not match those reported by Dahmann and Futcher [6], in which no significant effect on meiotic progress was observed in *clb4*Δ/*clb4*Δ mutants. We therefore tested whether this discrepancy is due to a difference in background strains - our evolution experiment used the *S. cerevisiae* Y55 strain which sporulates quickly and offers a strong selection against vegetative cells [7], while Dahmann and Futcher used W303. When we compared the *CLB4* deletion phenotype in the Y55 strain to other strains, we found that it is strain-specific: in two wild isolates, Y55 and its close relative SK1 [18], *clb4*Δ/*clb4*Δ diploids mostly form dyads, whereas in W303, a laboratory strain, the same genotype produces tetrads. This study reveals a previously unknown function of a conserved cyclin and emphasizes the importance of studying biological mechanisms in diverse natural isolates.

## Results

### Loss of Clb4 rescues spore viability in cells lacking recombination

Homolog segregation in meiosis I is directed by recombination, which depends on double-stranded breaks produced by Spo11 [15]. In *spo11*Δ/*spo11*Δ diploids, only 1% of spores receive at least 1 copy of each chromosome and are viable. We tried to evolve recombinationless meiosis by repeatedly sporulating a *spo11*Δ/*spo11*Δ Y55 diploid, killing any unsporulated cells by exposing them to diethyl ether [7], and germinating the survivors (Fig. 2A). High spore viability was recovered within 4-6 cycles of this selection (Fig. 2B). The predominant meiotic products in these populations were dyads of two diploid spores rather than tetrads of four haploid spores. Sequencing the evolved strains revealed that they contained single homozygous mutations. Four strains contained mutations in *SPO13* or *MAM1*, genes whose inactivation results in the separation of sister centromeres in a single meiotic division [12, 27, 29, 33, 35]. The other ten strains all contained mutations in *CLB4*; seven were nonsense mutations (Fig. 2C). This suggested that Clb4 inactivation rescues spore viability in *spo11*Δ/*spo11*Δ diploids and leads to dyad formation.

**Figure 2.**
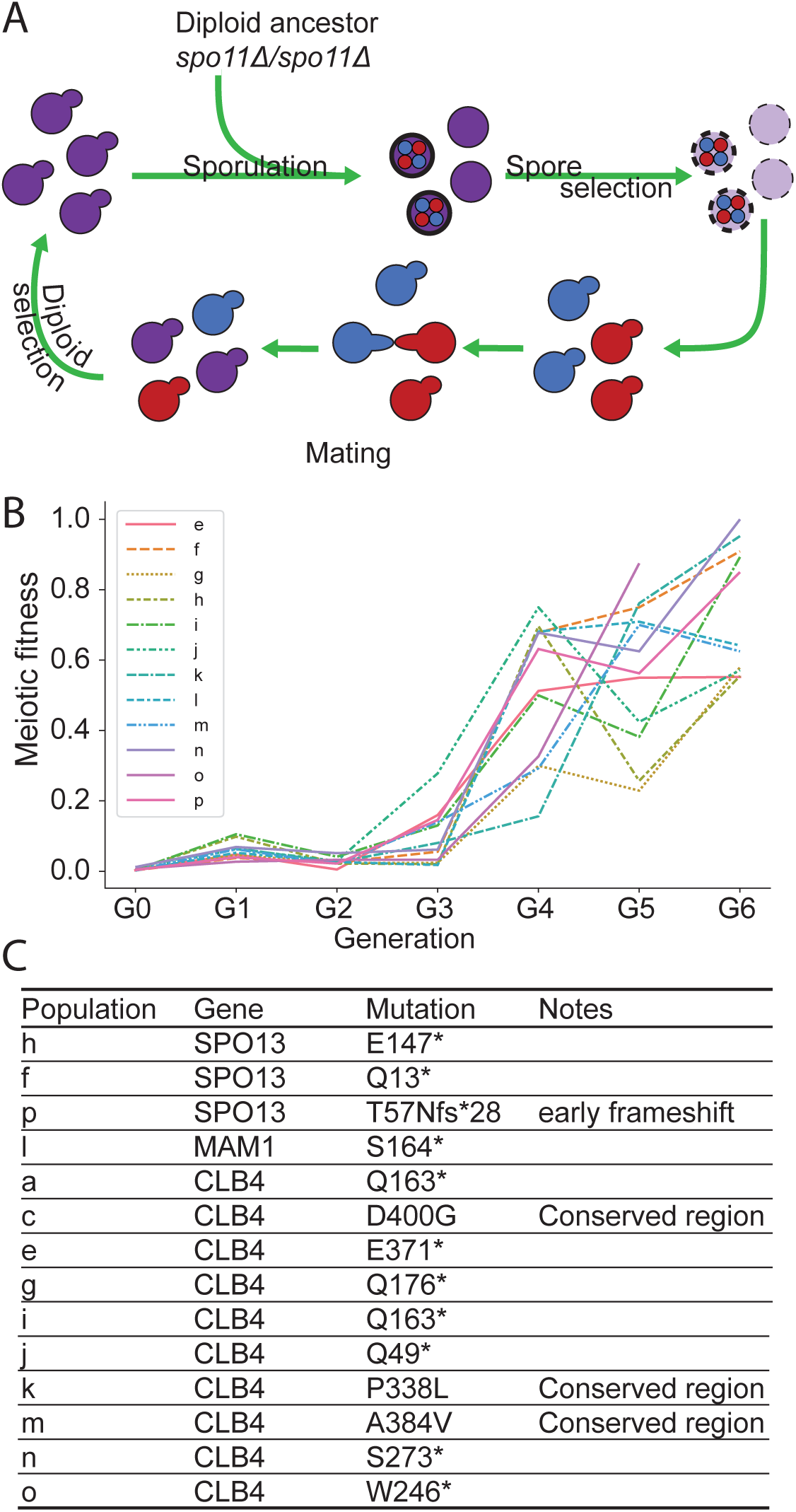
Experimental evolution of a Y55 *spo11*Δ strain. **(A)** Schematic of the experimental design. Y55 *spo11*Δ (yGL0057) was evolved in 16 populations across 2 experiments, one for populations a-d and another for populations e-p. Populations b and d were eliminated due to contamination. Each cycle is one sexual generation. See methods for detailed experimental evolution protocol. **(B)** Meiotic fitness across sexual generations in populations e-p. Meiotic fitness was measured as the ratio of colony forming units (CFU) after spore selection to CFU before spore selection. **(C)** Predominant mutations in each population. “*” notes stop codon. The listed mutations were identified in whole genome sequence analysis of the evolved populations after 6 selection cycles, i.e G6.

### Deletion of *CLB4* leads to dyad formation in the Y55 and SK1 strains, but not in W303

To confirm that inactivating *CLB4* leads to dyad formation, we constructed diploid *clb4*Δ/*clb4*Δ *spo11*Δ/*spo11*Δ strains. These strains formed dyads at a significantly higher rate than wild-type Y55 diploids (Fig. 3A). The double mutant *clb4*Δ/*clb4*Δ *spo11*Δ/*spo11*Δ diploids formed dyads at only a slightly higher rate than *clb4*Δ/*clb4*Δ, suggesting dyad formation is not dependent on the absence of Spo11. Since this result differs from that of Dahmann and Futcher [6], we also tested the phenotype of *clb4*Δ/*clb4*Δ in the W303 background. As previously observed, the W303 *clb4*Δ/*clb4*Δ diploids produced mostly tetrads. In contrast, *clb4*Δ/*clb4*Δ diploids of the SK1 strain, which is closely related to Y55 [18] and commonly used in yeast meiosis studies, produce mostly dyads, suggesting that a genetic difference between W303 and the other two strains alters the meiotic phenotype caused by removing Clb4.

**Figure 3.**
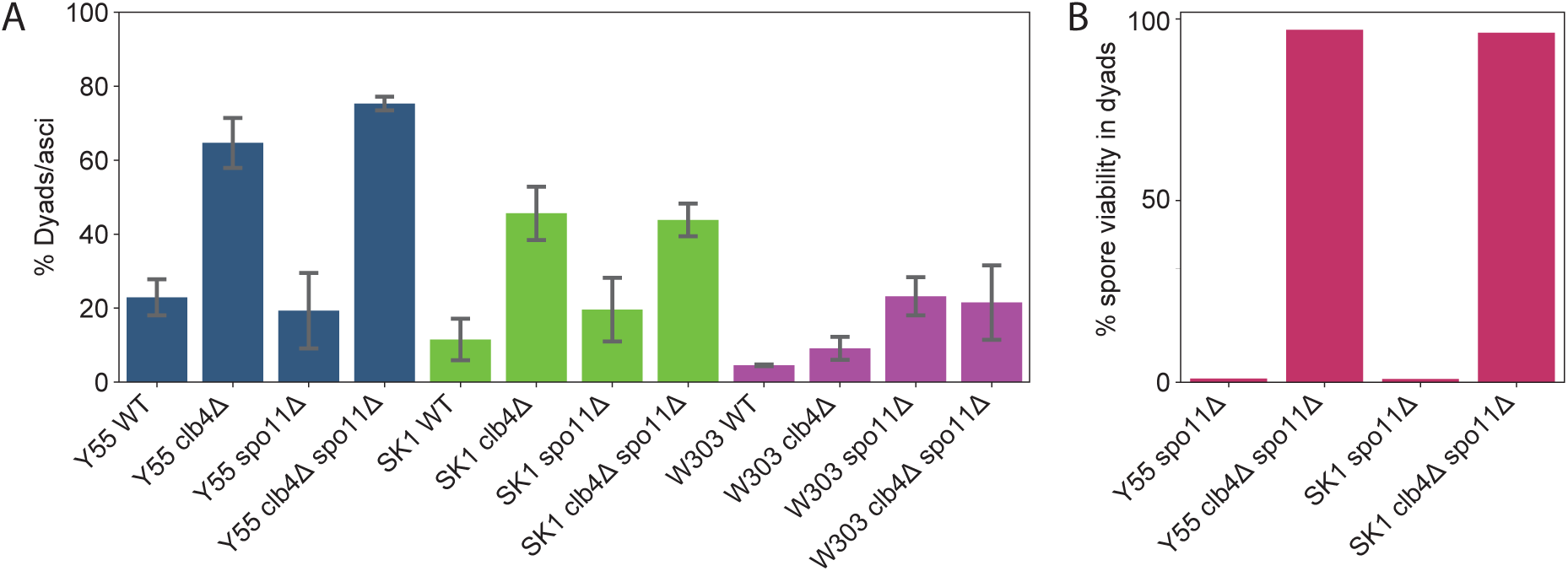
Phenotypic analysis of *CLB4* loss-of-function mutants. **(A)** Fraction of asci containing two spores in the following strains: Y55 wild-type (Y55 WT, yGL0340), Y55 *clb4*Δ (yGL0341), Y55 *spo11*Δ (yGL0342), Y55 *clb4*Δ *spo11*Δ (yGL0344), SK1 wild-type (SK1 WT, yGL0448), SK1 *clb4*Δ (yGL0449), SK1 *spo11*Δ (yGL0450), SK1 *clb4*Δ *spo11*Δ (yGL0451), W303 wild-type (W303 WT, yGL0123), W303 *clb4*Δ (yGL0160), W303 *spo11*Δ (yGL0003) and W303 *clb4*Δ *spo11*Δ (yGL0170). All listed mutations are homozygous. 3 replicates were made from each YPD culture during dilution into pre-sporulation media. The number of dyads and triads plus tetrads in each strain was compared between the following strains using a chi-square test. The tests were performed on pairs of replicates - the following are the tests with the highest p-value from each strain comparison: Y55 WT vs Y55 *clb4*Δ, *χ*^2^(*df* = 1*, N* = 163) = 24.00*, p <* 0.00001; Y55 *spo11*Δ vs Y55 *clb4*Δ *spo11*Δ, *χ*^2^(*df* = 1*, N* = 225) = 43.06*, p <* 0.00001; SK1 WT vs SK1 *clb4*Δ, *χ*^2^(*df* = 1*, N* = 344) = 11.24*, p* = 0.0008; SK1 *spo11*Δ vs SK1 *clb4*Δ *spo11*Δ, *χ*^2^(*df* = 1*, N* = 172) = 5.60*, p* = 0.01801; W303 WT vs W303 *clb4*Δ, *χ*^2^(*df* = 1*, N* = 223) = 0.00*, p* = 1; W303 *spo11*Δ vs W303 *clb4*Δ *spo11*Δ, *χ*^2^(*df* = 1*, N* = 197) = 0.45*, p* = 0.5. **(B)** The fraction of viable spores from dissected dyads in the Y55 and SK1 strains listed above. All listed mutations are homozygous. Dyads were collected from the same cultures used to generate the data in panel A.

In the absence of recombination, the viability of the spores from dyads reveals the pattern of centromere segregation in the division that produced them. If the sister centromeres failed to separate, homologs would segregate independently of each other, producing massive aneuploidy, and the vast majority of spores would be inviable. But if the sister centromeres separate and segregate from each other, the spores would be diploid, euploid, and viable. The high spore viability of Y55 *clb4*Δ/*clb4*Δ *spo11*Δ/*spo11*Δ dyads (Fig. 3B) suggested that Clb4, like Spo13 and Mam1, is required to prevent sister segregation from occurring in meiosis I.

### Deletion of *CLB4* causes sister separation in meiosis I

To more directly test whether deletion of *CLB4* causes sisters to separate in meiosis I in Y55, we used time-lapse microscopy to follow the behavior of a marked chromosome as cells went through meiosis. We used a GFP-lactose repressor fusion protein (GFP-LacI) to see a lactose operator (LacO) array at the LEU2 locus [32, 34], near the centromere of chromosome III and an mCherry-tagged spindle pole body protein, Spc42 [8], to see the two rounds of spindle pole body separation that mark the two meiotic divisions (Fig. 4A-F). Sisters separated in meiosis I in a significantly higher fraction of *clb4*Δ/*clb4*Δ and *clb4*Δ/*clb4*Δ *spo11*Δ/*spo11*Δ cells compared to wild type (Fig. 4G). To follow chromosome segregation in dyads genetically, we used strains that were heterozygous for the centromere-linked gene *LEU2*. If homologs segregate from each other in the single meiotic division, one spore will be Leu+ and the other will be Leu-, whereas if the sisters separate, both spores will be Leu+. In the Y55 *clb4*Δ/*clb4*Δ strain, 75% of the dyads with two viable spores produced 2 Leu+ spores and in the *clb4*Δ/*clb4*Δ *spo11*Δ/*spo11*Δ strain this fraction was even higher at 100% (Fig. 4H). Thus genetic and cell biological analysis support the conclusion that in Y55, Clb4 prevents premature sister separation in meiosis I. Although the fraction of divisions with sister separation is lower in cells that were followed by time lapse, the qualitative effect of deleting Clb4 is consistent between analyses by microscopy and genetics.

**Figure 4.**
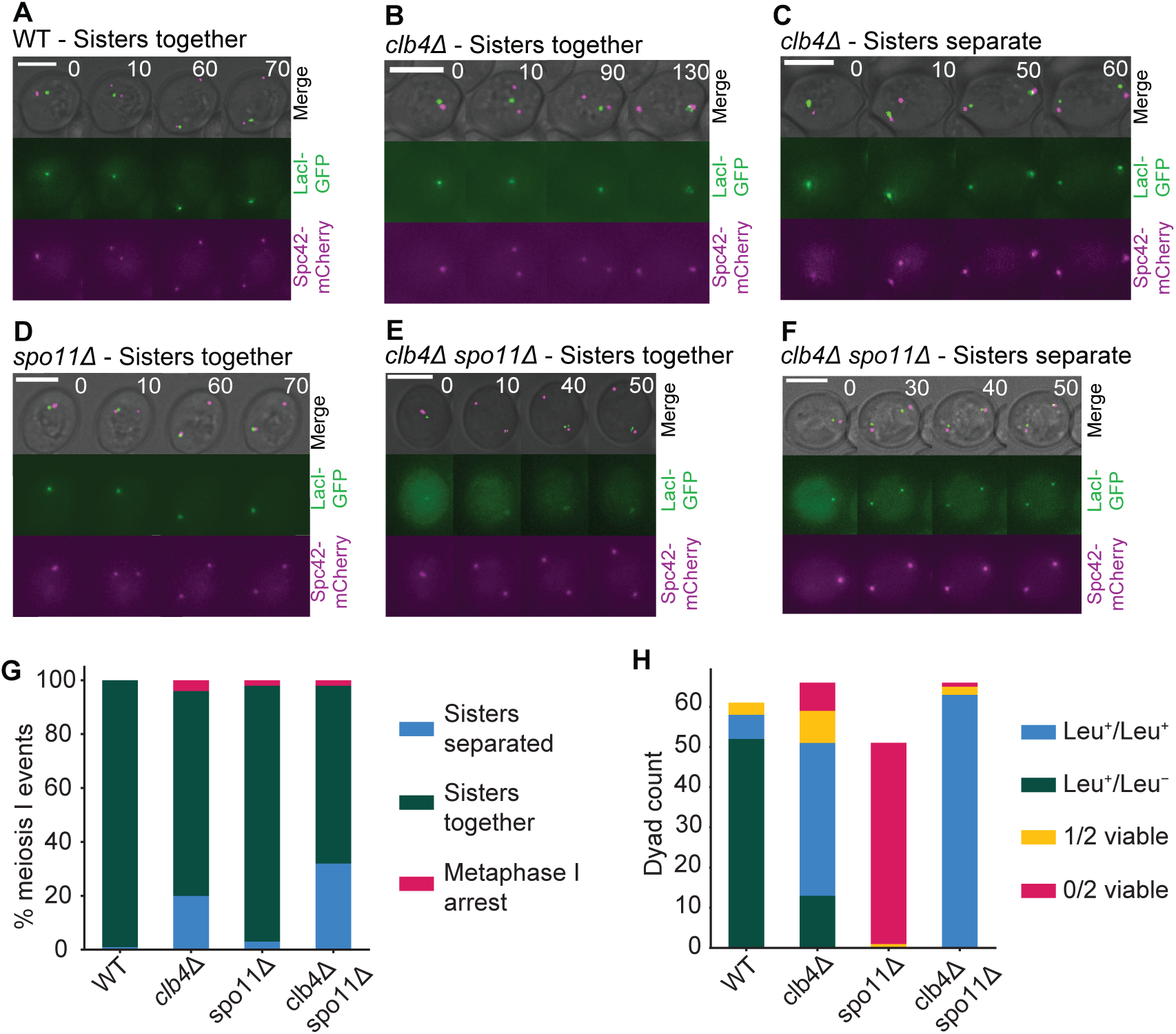
Sister separation in meiosis I in *CLB4* and *SPO11* mutants. **(A-F)** Representative images of meiosis I from time lapse movies of sporulation in the following Y55 strains: wild-type (WT, LY9542), *clb4*Δ (LY9361), *spo11*Δ (LY9464) and *clb4*Δ *spo11*Δ (LY9518). Cultures were incubated for 9 hours in bulk culture in sporulation medium before being transferred to a microscopy chamber. See methods for detailed protocol. Meiosis I events were identified by the first duplication of the spindle pole body (SPB) and subsequent SPB separation. **(G)** The fraction of meiosis I events observed in time lapse movies where sisters had separated vs. stayed together. 100 events were recorded and collected over 2 separate movies for each strain, taken on different days. The number of meiosis I sister separation events in each strain was compared to wild-type (WT) using a chi-square test: *clb4*Δ, *χ*^2^(*df* = 1*, N* = 196) = 20.14*, p <* 0.00001; *spo11*Δ, *χ*^2^(*df* = 1*, N* = 198) = 1.08*, p* = 0.30; *clb4*Δ *spo11*Δ, *χ*^2^(*df* = 1*, N* = 198) = 35.74*, p <* 0.00001. Additionally, the same analysis was made between *clb4*Δ and *clb4*Δ *spo11*Δ: *χ*^2^(*df* = 1*, N* = 194) = 3.4532*, p* = 0.06. **(H)** Spore viability and segregation pattern of dissected dyads. Strains used were Y55, homozygous for the listed deletions and LEU2 heterozygotes (*LEU2* /*leu2*Δ::*3xHA*. Sporulation was performed in bulk culture (see methods), and dyads were dissected on YPD plates, incubated for ∼24 hours and replica plated to CSM-leucine to asses leucine prototrophy. The number of Leu^+^/Leu^+^ and Leu^+^/Leu*^−^*in the following strains were compared to WT using a chi-square test: *clb4*Δ, *χ*^2^(*df* = 1*, N* = 109) = 46.41*, p <* 0.00001; *clb4*Δ *spo11*Δ, *χ*^2^(*df* = 1*, N* = 121) = 99.05*, p <* 0.00001.

Additionally, to test whether *clb4*Δ/*clb4*Δ dyads are the result of a single division, we tracked chromatin distribution throughout meiosis using a Histone H2B-GFP fusion protein (HTB2-GFP). By starting observation of the sporulated cultures later than in the experiment in Figure 4, we were able to follow meiosis to the point of spore formation. Out of 72 *clb4*Δ/*clb4*Δ cells that formed dyads, 67 went through a single division of their chromatin content (Fig. S1G).

To perform a more detailed analysis of meiosis in *clb4*Δ/*clb4*Δ SK1 cells, we adopted a protocol that has been used in SK1 to examine the effects of altering the timing of gene expression, including that of individual cyclin genes. We regulated expression of Ndt80, the transcription factor that induces the meiotic divisions (Ndt80 block-release synchronization) [24]. *NDT80* was placed under the control of a beta-estradiol inducible promoter that was activated 4.5 hours after transferring SK1 *clb4*Δ/*clb4*Δ cells to sporulation medium. To track sister separation, we used a GFP-tet repressor fusion protein (GFP-tetR) to see a heterozygous tet operator (tetO) array [23] integrated near the centromere of chromosome V (CENV-GFP dots). Cells were collected 3 hours after *NDT80* induction and fixed, then stained with DAPI to visualize the nuclei. We analyzed CENV-GFP dot localization alongside the divisions of the nuclei to assess sister separation (Fig. 5A). Compared to the wild-type, the *clb4*Δ/*clb4*Δ strain had a significantly higher fraction of cells with 2 nuclei where each nucleus had a single GFP dot, indicating sister centromere separation in the first meiotic division. The same cultures were also sampled after 24 hours to assay their dyad fraction, as well as their viability and genetic segregation pattern (using the *HIS3* marker linked to the CENV-tetO array) of spores derived from dyads (Fig. 5B-D). These results are consistent with those from the Y55 strain, indicating that Clb4 has a similar role in both strains and that perturbing the normal sequence of meiotic gene expression did not influence the phenotype of removing Clb4.

**Figure 5.**
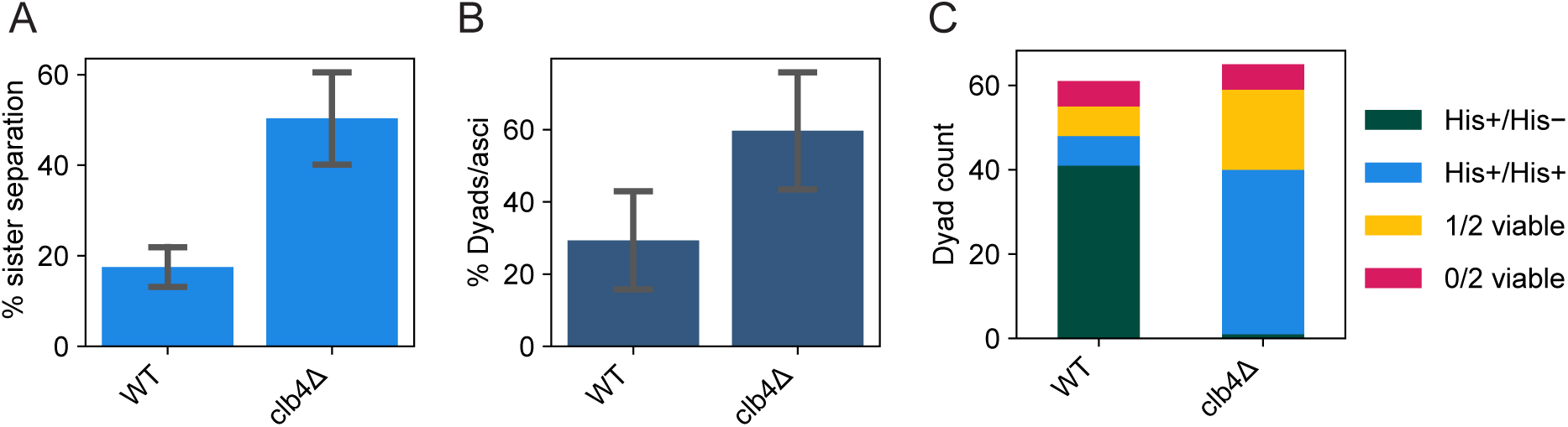
Meiotic time course and phenotypic characterization of a synchronized SK1 *clb4*Δ/*clb4*Δ strain. **(A)** Fraction of cells with two nuclei (binucleates) and one GFP dot in each nucleus, indicating meiosis I sister centromere separation, in the following strains: wild-type (WT, yGL0477) and *clb4*Δ (yGL0475), 3 hours after *NDT80* induction. The *CLB4* deletion is homozygous. Data was collected across 3 replicates, from time courses performed on separate days. The number of binucleates that separated sister centromeres was compared between WT and *clb4*Δ using a chi-square test. The tests were performed on pairs of replicates - the test with the highest p-value was: *χ*^2^(*df* = 1*, N* = 200) = 7.95*, p* = 0.00481. **(B)** Dyad fraction in the strains listed for panel A, in the mature (*>* 24 hour old) sporulation cultures. Data was collected across 3 replicates. The number of dyads in *clb4*Δ was compared to wild-type using a chi-square test. The tests were performed on pairs of replicates - the test with the highest p-value was: WT vs *clb4*Δ, *χ*^2^(*df* = 1*, N* = 200) = 8.18*, p* = 0.00423. **(C)** Number of viable spores per dyad and the segregation pattern of a CENV-*HIS3* marker in dyads with 2 viable spores, dissected from dyads in the strains listed for panel A. Dyads were collected from the same cultures used for panels A and B. The number of His^+^/His^+^ and His^+^/His*^−^* dyads in *clb4*Δ was compared to wild-type using a chi-square test: *χ*^2^(*df* = 1*, N* = 88) = 56.85*, p <* 0.00001.

### Dyad formation in the absence of Clb4 is not dependent on the spindle checkpoint

The *CLB4* deletion phenotype is similar to that of deleting *SPO13*. Previous work demonstrated that dyad formation in *spo13*Δ/*spo13*Δ is dependent on the spindle checkpoint [33]. We asked whether the same is true for *clb4*Δ/*clb4*Δ. We deleted *MAD2*, a component of the spindle checkpoint complex [5, 17], in a Y55 *clb4*Δ strain and a Y55 *spo13*Δ strain, and assessed the frequency of dyad formation in homozygous diploids (Fig. 6). The absence of Mad2 had significant effects on both *clb4*Δ/*clb4*Δ and *spo13*Δ/*spo13*Δ, but in opposite directions. The *spo13*Δ/*spo13*Δ *mad2*Δ/*mad2*Δ dyad fraction was significantly smaller compared to *spo13*Δ/*spo13*Δ, close to the dyad fraction of the wild-type. However, the dyad fraction of *clb4*Δ/*clb4*Δ *mad2*Δ/*mad2*Δ was significantly larger than both wild-type and *clb4*Δ/*clb4*Δ. This suggests that the termination of meiosis after a single division is caused by different mechanisms in Y55 *clb4*Δ/*clb4*Δ and *spo13*Δ/*spo13*Δ.

**Figure 6.**
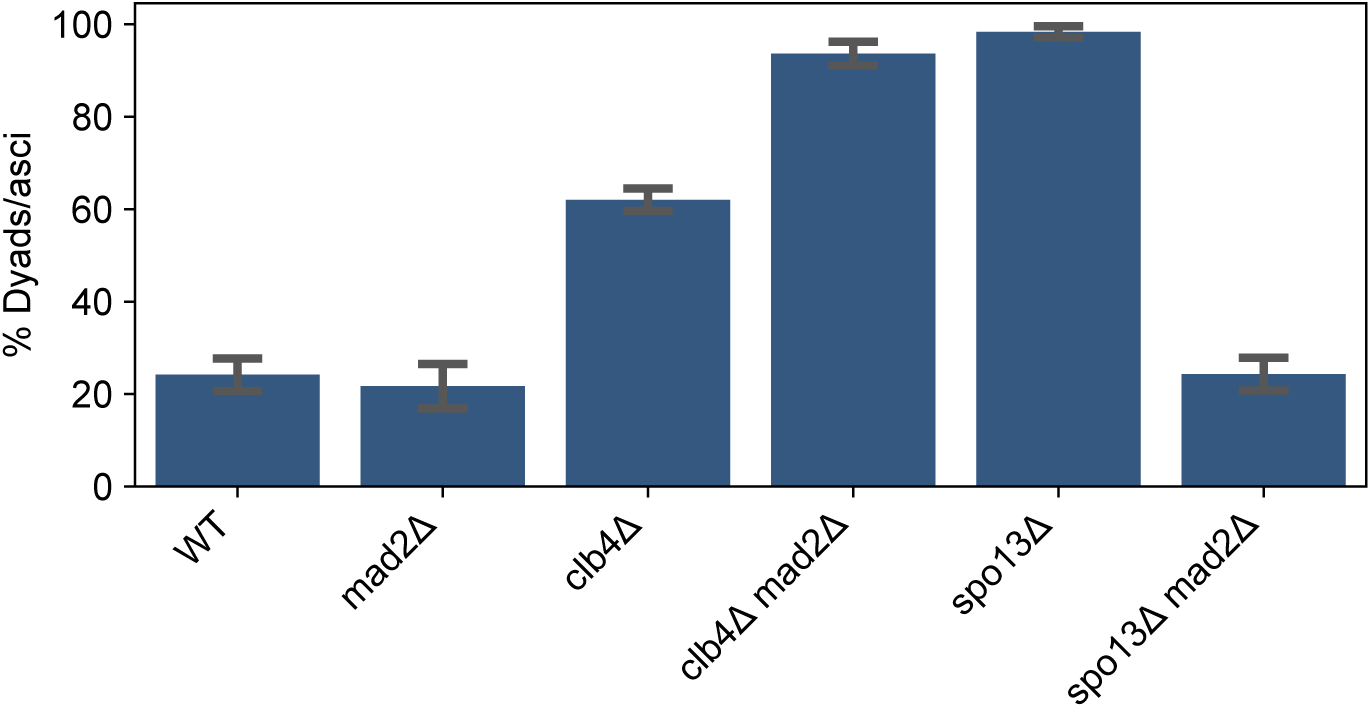
The effects of the spindle checkpoint on the fraction of dyads in *CLB4* and *SPO13* mutants. The fraction of two-spored asci (dyads) in the following strains: wild-type (WT, yGL0340), *mad2*Δ (yGL0349), *clb4*Δ (yGL0341), *clb4*Δ *mad2*Δ (yGL0350), *spo13*Δ (yGL0492), *spo13*Δ *mad2*Δ (yGL0447). All listed mutations are homozygous. Data for Y55 WT and *clb4*Δ is repeated from fig. 3A. 3 replicates were split from each YPD culture during dilution into pre-sporulation media. The number of dyads and tri/tetrads in each strain was compared between the following strains using a chi-square test. The tests were performed on pairs of replicates - the following are the tests with the highest p-value from each strain comparison: WT vs *mad2*Δ, *χ*^2^(*df* = 1*, N* = 231) = 0.38*, p* = 0.53878; WT vs *spo13*Δ, *χ*^2^(*df* = 1*, N* = 221) = 104.43*, p <* 0.00001; WT vs *spo13*Δ *mad2*Δ, *χ*^2^(*df* = 1*, N* = 244) = 0.36*, p* = 0.55039; WT vs *clb4*Δ *mad2*Δ, *χ*^2^(*df* = 1*, N* = 236) = 94.03*, p <* 0.00001; *spo13*Δ vs *spo13*Δ *mad2*Δ, *χ*^2^(*df* = 1*, N* = 223) = 112.30*, p <* 0.00001; *clb4*Δ vs *clb4*Δ *mad2*Δ, *χ*^2^(*df* = 1*, N* = 252) = 23.42*, p <* 0.00001;

### Clb1 is required for meiosis I sister separation in the absence of Clb4

We next sought to reexamine other *CLB* mutants and analyze their genetic interactions with *spo11*Δ. Previous work reported that the double mutant *clb1*Δ/*clb1*Δ *clb4*Δ/*clb4*Δ goes through a single meiotic division, in which homologs are segregated rather than sisters, in both the W303 [6] and SK1 [14] strain backgrounds. We constructed Y55 strains with different combinations of homozygous deletions in *CLB1*, *CLB3* and *CLB4*. Each strain also had two heterozygous centromere-linked markers (*LEU2* /*leu2*Δ and *TRP1* /*trp1*Δ), allowing us to determine whether homologs or sister centromeres had segregated in the single division that produced two viable spores. We measured the frequency of dyad formation (Fig. 7, top) and dissected dyads from each strain to test their spore viability and segregation pattern (Fig. 7, middle and bottom panels).

**Figure 7.**
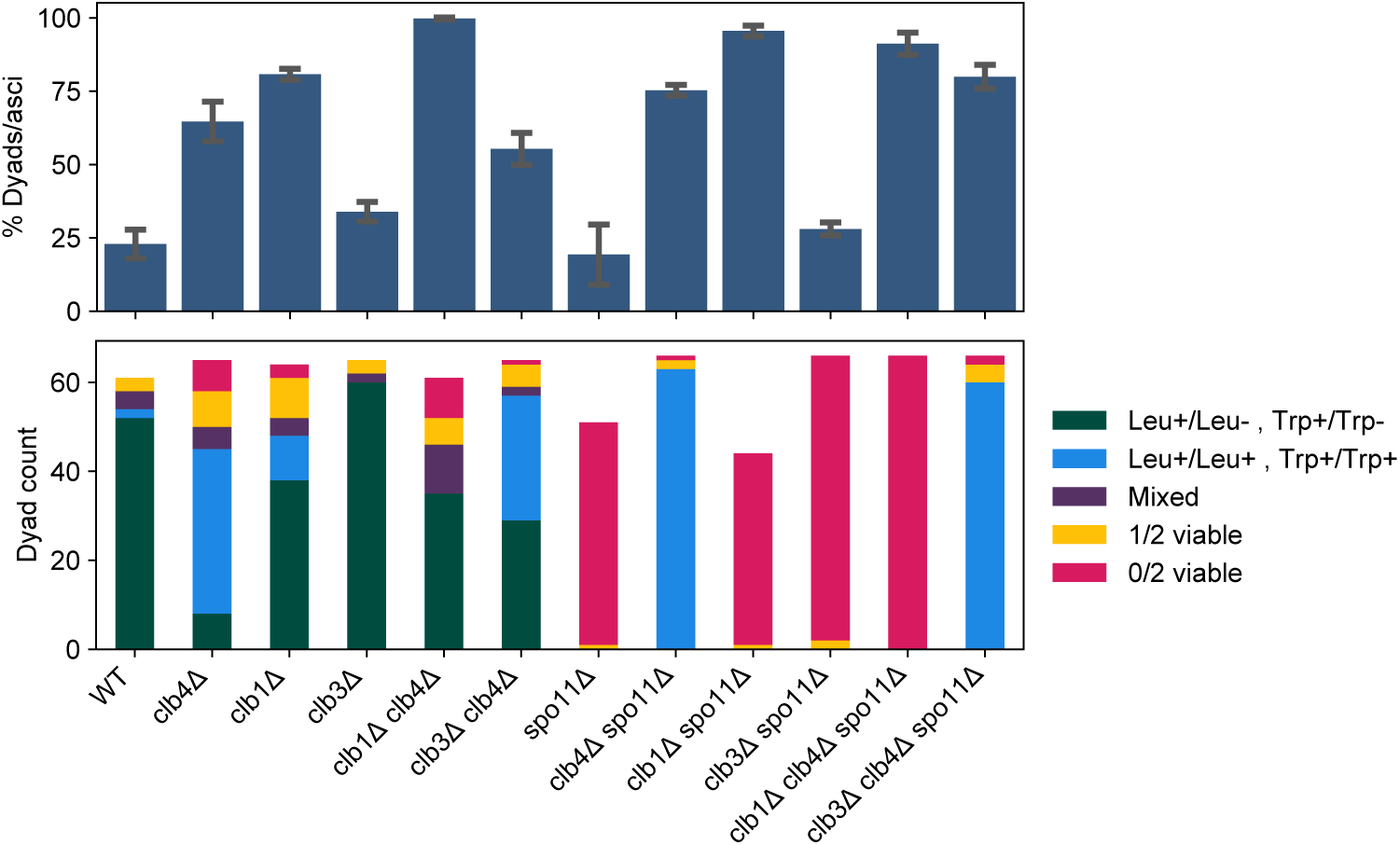
Dyad fraction, viability and segregation pattern in *CLB* and *SPO11* mutants. Mean dyad fraction (top) and number of viable spores per dyad and the segregation pattern of *LEU2* and *TRP1* markers (bottom) in the following strains: wild-type (WT, yGL0340), *clb4*Δ (yGL0341), *clb1*Δ (yGL0388), *clb3*Δ (yGL0343), *clb1*Δ *clb4*Δ (yGL0389), *clb3*Δ *clb4*Δ (yGL0345), *spo11*Δ (yGL0342), *clb4*Δ *spo11*Δ (yGL0344), *clb1*Δ *spo11*Δ (yGL0390), *clb3*Δ *spo11*Δ (yGL0346), *clb1*Δ *clb4*Δ *spo11*Δ (yGL0391) and *clb3*Δ *clb4*Δ *spo11*Δ (yGL0347). All strains are from the Y55 background and homozygous for the listed deletions, as well as heterozygous for *LEU2* and *TRP1*. Data for Y55 WT and *clb4*Δ is repeated from figure 3A. Three replicates were made from each YPD culture during dilution into pre-sporulation media. The number of dyads and tri/tetrads in each strain was compared between the following strains using a chi-square test. The tests were performed on pairs of replicates - the following are the tests with the highest p-value from each strain comparison: WT vs *clb1*Δ, *χ*^2^(*df* = 1*, N* = 246) = 56.83*, p <* 0.00001; WT vs *clb3*Δ, *χ*^2^(*df* = 1*, N* = 210) = 1.53*, p* = 0.21639; *clb4*Δ vs *clb1*Δ *clb4*Δ, *χ*^2^(*df* = 1*, N* = 208) = 24.51*, p <* 0.00001; *clb4*Δ vs *clb3*Δ *clb4*Δ, *χ*^2^(*df* = 1*, N* = 205) = 0.06*, p* = 0.80898; *spo11*Δ vs *clb1*Δ *spo11*Δ, *χ*^2^(*df* = 1*, N* = 216) = 80.57*, p <* 0.00001; *spo11*Δ vs *clb3*Δ *spo11*Δ, *χ*^2^(*df* = 1*, N* = 277) = 0.09*, p* = 0.76870; *clb4*Δ *spo11*Δ vs *clb1*Δ *clb4*Δ *spo11*Δ, *χ*^2^(*df* = 1*, N* = 200) = 2.01*, p* = 0.15580; *clb4*Δ *spo11*Δ vs *clb3*Δ *clb4*Δ *spo11*Δ, *χ*^2^(*df* = 1*, N* = 208) = 0.01*, p* = 0.93618. ∼22 spores were dissected per replicate and pooled for analysis. Viability and *LEU2* segregation data is repeated from figure 4H. The number of Leu^+^/Leu^+^ Trp^+^/Trp^+^ and Leu^+^/Leu*^−^* Trp^+^/Trp*^−^* in the following strains were compared to WT using a chi-square test: WT vs *clb4*Δ, *χ*^2^(*df* = 1*, N* = 99) = 63.38*, p <* 0.00001; WT vs *clb1*Δ, *χ*^2^(*df* = 1*, N* = 102) = 7.18*, p <* 0.008; WT vs *clb3*Δ, *χ*^2^(*df* = 1*, N* = 114) = 2.26*, p* = 0.13; *clb4*Δ vs *clb1*Δ *clb4*Δ, *χ*^2^(*df* = 1*, N* = 80) = 53.54*, p <* 0.00001; *clb4*Δ vs *clb3*Δ *clb4*Δ, *χ*^2^(*df* = 1*, N* = 102) = 11.92*, p* = 0.00056.

As previously observed in other strain backgrounds, Y55 *clb1*Δ/*clb1*Δ *clb4*Δ/*clb4*Δ diploids formed dyads almost exclusively. Out of the 48 dyads that contained two viable spores, 10 contained two Leu+ spores while 38 contained only one Leu+ spore. For *TRP1*, the pattern was even more pronounced, with only one of 48 dyads containing 2 Trp+ spores, supporting the conclusion that the activity of Clb1 is required for meiosis I sister separation in cells lacking Clb4.

While the *clb1*Δ/*clb1*Δ *clb4*Δ/*clb4*Δ phenotype is consistent across strains, Y55 *clb1*Δ/*clb1*Δ produces ∼80% dyads, unlike W303 *clb1*Δ/*clb1*Δ which produces only 14% dyads [6]. In contrast to Y55 *clb4*Δ/*clb4*Δ dyads, centromere linked markers suggest that only ∼22% of Y55 *clb1*Δ/*clb1*Δ dyads segregated sister centromeres in meiosis I. This suggests that like Clb4, Clb1 is required for meiotic progression in Y55, but does not share Clb4’s role in preventing sister centromere separation in meiosis I.

As predicted by the low activity of Clb3 in meiosis I [4], deletion of *CLB3* did not affect the *clb4*Δ phenotype as significantly as *CLB1*.

### *clb1*Δ and *clb3*Δ are unable to rescue *spo11*Δ **spore viability**

We asked whether the deletion of other M-phase cyclins could cause sister centromere separation in meiosis I. The ability of precocious *CLB1* or *CLB3* expression to induce sister centromere separation [24] and the near absence of Clb3-associated Cdk1 activity in meiosis I [4] predict that deletion of either of these cyclin genes would not cause sister centromere separation in meiosis I. Dyads from both *clb1*Δ/*clb1*Δ and *clb3*Δ/*clb3*Δ diploids showed a similar frequency of sister separation in meiosis I as wild-type dyads (Fig. 7, bottom). In addition, and in contrast to *clb4*Δ/*clb4*Δ *spo11*Δ/*spo11*Δ dyads, dyads from both the *clb1*Δ/*clb1*Δ *spo11*Δ/*spo11*Δ and *clb3*Δ/*clb3*Δ *spo11*Δ/*spo11*Δ strains were almost exclusively inviable. Thus Clb4 is unique among M-phase cyclins in preventing sister centromere separation in meiosis I.

## Discussion

What we had intended as a route to experimentally evolve recombination-less meiosis in budding yeast turned out to be a powerful selection for mutations that cause sister centromere segregation in a single meiotic division. In addition to recovering mutations in genes of the monopolin complex, which prevents sister centromere segregation and separation in meiosis I, we repeatedly recovered mutations that inactivated the Clb4 cyclin. The phenotype of diploids lacking a functional copy of *CLB4* depends on the strain background. In two related strains, Y55 and SK1, Clb4 inactivation rescues the spore viability of *spo11*Δ/*spo11*Δ diploids by terminating meiosis after a single division, in which sister centromeres rather than homologous chromosomes are segregated. In the laboratory strain, W303, diploids lacking Clb4 produce tetrads. Sister centromere separation in *clb4*Δ/*clb4*Δ is dependent on the presence of Clb1 - *clb1*Δ/*clb1*Δ *clb4*Δ/*clb4*Δ cells produce dyads after segregating homologous chromosomes (Fig. 7), a phenotype that is consistent across the yeast strains used here [6, 14]. This suggests that the balancing activity of different cyclins is important for regulating meiotic chromosome segregation.

We suggest that the phenotypes of these cyclin mutants can be explained by effects on the timing of various events promoted by Cdk1 in combination with a timer that triggers spore formation after a fixed interval from the beginning of the first meiotic division. In this model, Clb4 accelerates the co-orientation of sister centromeres and the protection of cohesin near the centromeres, whereas Clb1 accelerates spindle assembly [24]. In the absence of Clb4, the fusion of sister centromeres into a single functional unit is delayed but spindle assembly is not, leading to sister separation in meiosis I and dyad formation. In the absence of Clb1, the spindle forms later and sister centromere behavior is normal, leading to homolog segregation in meiosis I, with the delay preventing a second division before the spore formation timer has expired (in this context, it is worth noting the high fraction of dyads in the absence of Clb1, with most of these dyads segregating homologs according to genetic marker data - see figure 7). In the absence of Clb1 and Clb4, the delay in spindle assembly allows enough time for a backup process to convert the sister centromeres into a single functional unit thus preventing sister centromere segregation in meiosis I. This timing model also offers an explanation of the differences between the effects of mutations in Y55 compared to W303. If the spore assembly timer runs faster in Y55, it is more likely that slowing down individual steps in meiosis will lead to dyad formation: even if removing Clb1 delays spindle assembly equally in Y55 and W303, the faster spore formation timer in Y55 will terminate meiosis before a second division has time to occur.

The model requires that some aspect of meiosis I is delayed in cells lacking Clb4 and the lack of a second division in *clb4*Δ/*clb4*Δ *mad2*Δ/*mad2*Δ says that unlike meiosis in *spo13*Δ/*spo13*Δ, the delay is not due to activation of the spindle checkpoint. In *spo13*Δ/*spo13*Δ diploids, sister centromeres attach to opposite poles, pericentromeric cohesin is not protected, and sisters separate at the same time that homologs would separate in a wild-type meiosis I. In monopolin mutants, sister centromeres also attach to opposite poles, but pericentromeric cohesin is still protected, and thus sister centromeres separate at the same time that they would separate in a wild-type meiosis II [35]. Because inactivating the spindle checkpoint does not lead to a second meiotic division in Y55 *clb4*Δ/*clb4*Δ diploids, we argue that removing Clb4 produces a phenotype that is more similar to that of monopolin mutants.

Do the small fraction of dyads found in wild type Y55 diploids also reflect sister centromere separation in meiosis I? We argue that they do not. Although Y55 meiosis does produce dyads in approximately 20% of cases where Clb4 is present, our results show these dyads to be qualitatively different from those produced in the absence of Clb4, in three main ways - First, time lapse microscopy reveals that *clb4*Δ/*clb4*Δ dyads are produced after a single division, whereas wild-type dyads undergo two divisions and end up “losing” two chromatin masses that do not get packed into a spore (Fig. S1A). Second, the vast majority of spores from Y55 *spo11*Δ/*spo11*Δ dyads are inviable (Fig. 4H). Third, the segregation pattern of genetic markers in Y55 wild-type dyads is predominantly consistent with homolog segregation and not sister centromere segregation (Fig. 7).

Our results reveal a strain difference in the requirement of Clb4 for proper meiotic progression, which raises the question of how widespread the Clb4 requirement is among *S. cerevisiae* strains. W303, which does not require Clb4, is a domesticated strain, selected to have properties useful for yeast geneticists [31], while the Y55 strain used in our evolution experiment remained largely unchanged since its isolation from wine grapes. This raises the possibility that requiring Clb4 for proper meiotic progression is the ancestral state of yeast meiosis, though studying additional strains from multiple lineages would be required before drawing any conclusions. Regardless, this difference highlights the importance of using multiple natural isolates in genetic studies.

Finally, the *spo11*Δ/*spo11*Δ evolution experiment we have designed, or selection screens similar to it, could be substantially scaled up to include many more populations, and used in the future to identify additional cellular components involved in ensuring sister centromere co-segregation in meiosis I.

## Acknowledgements

We thank the members of the Murray lab and Lacefield lab for helpful discussions, Sue Biggins, Mike Laub, Adele Marston and Matt Miller for comments on the manuscript, and Eļcin Ünal for strains, protocols and a helpful discussion. GL and AWM are supported by the grant NIH/NIGMS R01 GM043987. AWM is also supported by the NSF-Simons Center for the Mathematical and Statistical Analysis of Biology (NSF #1764269, Simons #594596). GL was also supported by a PhD fellowship from the Boehringer Ingelheim Fonds. SL and GC were supported by an NSF grant 2044556 and by core support through P20-GM113132 to SL.

## Materials and Methods

### Yeast strains

Yeast strains used in the experiments presented in this study came from either the Y55, W303 or SK1 backgrounds. All Y55 strains were derived from NKY177 (*MATa/MATα lys2*Δ/*lys2*Δ *HO/hoΔ::LYS2*), a gift from Nancy Kleckner. W303 strains were from a modified W303 background with replacement of the *BUD4* allele with its respective, functional allele from the S288C background. SK1 strains were gifts from Nancy Kleckner, Angelika Amon (through Xiaoxue Snow Zhou or Juliet Barker) or Elçin Ünal or derived from said gifted strains. Lists of all strains used, along with their genotypes, are available in supplemental file S1.

### Sporulation protocols

The following sporulation protocol was used in the experiments presented in figures 2, 3, 4G-H, 6 and 7. Cells were inoculated into culture tubes with rich medium, containing 2% yeast extract, 1% peptone and 2% dextrose (YPD). The YPD culture tubes were incubated in a roller drum at 30°C for ∼24 hours. YPD cultures were then diluted 1:50 into a pre-sporulation medium containing 2% yeast extract, 1% peptone and 1% potassium acetate (YPA). The YPA culture tubes were incubated in a roller drum at 30°C for 16-20 hours, while allowing gas exchange. YPA cultures were then pelleted (centrifuged) at 700g for 3 minutes. The supernatent was discarded and the pellet resuspended in sterile water (*>* 2-fold culture volume), pelleted again and resuspended in sporulation medium, containing 1% potassium acetate (1% KAc). The sporulation cultures tubes were incubated in a roller drum for *>* 48 hours, at either 30°C (for Y55) or 27°C (for W303). Culture volume was 2mL. SK1 strains were sporulated with a separate protocol. For the experiments presented in figures 3, sporulation was carried out as described above, with the only difference being the use of baffled flasks incubated in a shaker (at 30°C) in place of tubes. Culture volume was one tenth the maximum volume of the flasks.

### Experimental Evolution

The following protocol was used for each cycle of the evolution experiment presented in figure 2. Diploid cells were put through the sporulation protocol detailed above. Vegetative cells were then killed by treating the culture with diethyl ether. 0.5mL of sporulation culture and 0.5mL of diethyl ether were added to a 1.5mL microcentrifuge tube. The tubes were vortexed for 30 seconds at maximum speed on a benchtop vortex. After vortexing, the mixture quickly separates into 2 phases, with a lower aqueous phase containing the treated culture. The aqueous phase was removed and inoculated into 2mL YPD in a fresh 15mL culture tube (this marked the beginning of a new generation or cycle of the experiment). The YPD culture tubes were incubated in a roller drum at 30°C for 16-24 hours, after which they were taken out of the roller drum and incubated without agitation at 30°C for an additional 5-8 hours, to allow mating of haploid cells. Next, 20µL of each culture were inoculated into YPD medium containing 100µg/mL of ClonNat (Nourseothricin) and 300µg/mL of Hygromycin B ”(selecting for resistance markers inserted next to the MAT locus), to select for diploids. The diploid-selective YPD cultures were then incubated in a roller drum at 30°C for ∼24 hours, serving as the first step of the sporulation protocol and followed by dilution into YPA as detailed above. In the first cycle, cells from the yGL0057 ancestor (Y55, *spo11*Δ/*spo11*Δ) were entered into the sporulation protocol instead of diploid-selective culture.

### Ascus dissection protocol

The following protocol was used to dissect asci (dyads or tetrads) from sporulation cultures. A sample of sporulation culture with approximately 10^8^ cells/mL was mixed 9:1 with a 5U/µL solution of Zymolyase (Zymo Research, cat. E1004) to a final zymolyase concentration of 0.5U/µL and incubated at 30°C for 5 minutes. The mixture was diluted 1:9 in sterile water and stored in 4°C until used. 10µL from the diluted mixture were spread in a line on a YPD agar plate (2% agar, 2% yeast extract, 1% peptone and 2% dextrose) and allowed to dry. Asci on the YPD plate were dissected using a microscope equipped with a micromanipulator.

### Sporulation time course protocol

For the time course experiment presented in figures 5, sporulation was carried out with the following protocol, which is a slightly modified version of a protocol received from Elçin Ünal - Cells were patched onto solid media containing 2% agar, 2% yeast extract, 1% peptone and 3% glycerol (YPG). The YPG plate was incubated overnight (approximately 16 hours) at 30°C. Cells from the YPG plate were then patched onto a 4% YPD agar (2% agar, 2% yeast extract, 1% peptone and 4% dextrose) plate, and incubated at 30°C for 7-8 hours. Cells from the 4% YPD plate were then used to inoculate a liquid YPD (2%) culture in a baffled flask, which was incubated in a shaker (at *>* 300rpm) for ∼24 hours at room temperature to reach O.D_600_ *>* 10. The YPD culture was diluted to O.D_600_ = 0.25 into a pre-sporulation medium containing 1% yeast extract, 2% tryptone, 1% potassium acetate and 50mM potassium phthalate (BYTA), in baffled flasks, and incubated in a shaker for 15-16 hours at 30°C to reach O.D_600_ *>* 5. BYTA pre-sporualtion cultures were then transfered to conical tubes and pelleted at 700g for 3 minutes. The supernatent was discarded and the pellet resuspended in sterile water (*>* 2-fold culture volume), pelleted again and resuspended in sporulation medium containing 2% potassium acetate and 0.02% raffinose (SPO). Sporulation cultures were then transferred to baffled flasks and incubated in a shaker at 30°C. Culture volume was one tenth the maximum volume of the flasks.

*NDT80* expression was induced by adding β-estradiol (10mM in ethanol stock solution) to a final concentration of 1µM, 4 hours and 30 minutes after transfer to sporulation medium. Samples were taken from each culture and fixed with the fixation protocol below. Assays of the mature sporulation cultures (sporulation rate, dyad fraction and dissections) were performed *>* 24 hours after transfer to sporulation medium.

### Fixation protocol

The following fixation protocol was used to fix cells going through meiosis in the time course experiments presented in figure 5. It was received from Elçin Ünal and used by Miller, Ünal, Brar and Amon, 2012 [24]. A 500µL sample was taken from each culture and placed in a microcentrifuge tube. 55µL of 37% formaldehyde was added to each sample to a final concentration of 3.7%. Samples were briefly vortexed and incubated at room temperature for 15 minutes. Samples were then pelleted at 9,400g for 30 seconds. The supernatent was then aspirated and the cells were resuspended in a 100µL of a solution containing 83.4mM K_2_HPO_4_, 16.6mM KH_2_PO_4_, 1.2M sorbitol and 1% Triton X-100, and incubated at room temperature for 5 minutes for permeabilization. Samples were then pelleted at 9,400g for 30 seconds. The supernatent was then aspirated and the cells were resuspended in a 50µL of a solution containing 83.4mM K_2_HPO_4_, 16.6mM KH_2_PO_4_, 1.2M sorbitol. The fixed cells’ DNA was stained by adding 4’,6-diamidino-2-phenylindole (DAPI) to a final concentration of 0.05µg/mL. The fixed samples were stored in 4°C for up to a month before imaging.

### DAPI and GFP-dot imaging protocol

The following protocol was used to image fixed cells in the time course experiments presented in figure 5. Imaging was performed using a Nikon Ti inverted microscope, with a 100x oil-immersion objective (NA = 1.4), on glass slides treated with ConA (2 mg/mL concanavalin A, 5mM MnCl_2_, 5mM CaCl_2_). For each field of view, a Z series of 11 images were taken in the GFP and DAPI channels, separated by 0.3µm, and a single brightfield image was taken at the center of the Z series. Images were processed in the Fiji distribution of ImageJ (National Institutes of Health). DAPI and GFP Z series were processed by maximum intensity projection, and contrast was enhanced to allow better viewing.

### Time lapse movies of sporulation

The following protocol was used for time lapse microscopy experiments (movies), data from which was included in figures 4 and S1. Cells were grown in 2x synthetic complete medium (2xSC; 0.67% bacto–yeast nitrogen base without amino acids, 0.2% dropout mix with all amino acids, and 2% glucose) overnight at 30*^◦^*C, transferred to 2xSCA (1:40) (0.67% bacto–yeast nitrogen base without amino acids, 0.2% dropout mix with all amino acids, 1% potassium acetate) at 30*^◦^*C for 12-16 hours, washed with water twice, and transferred to 1% potassium acetate at 25*^◦^*C for 9 hours. All movies were taken in a chamber on a coverslip coated with Concanavalin A (1mg/mL; Sigma; used 1xPBS as solvent). Prior to imaging, cells were concentrated and loaded into the coverslip in a chamber. An agar pad (0.05g/mL) containing 1% potassium acetate was used to make a monolayer of cells and removed before imaging. Pre-conditioned 1% potassium acetate was added to the chamber before imaging. All meiosis movies were run for 12hrs. LacO:TRP1 and pCUP1-LacI-GFP:HIS3 expressing cells were imaged at room temperature using a DeltaVision pDV microscope (Applied Precision) equipped with a CoolSNAP HQ2/HQ2-ICX285 camera using a 60-100x oil objective (U-Plan S-Apochromat-N, 1.4 NA). During time-lapse imaging, five Z steps (0.8µm) were acquired every 10 min. The exposure times used for brightfield, LacI-GFP, and Spc42-mCherry were 0.3ms, 0.1ms, and 0.25ms, respectively, with neutral-density filters transmitting 10–32% of light intensity. Images were acquired using SoftWoRx software (GE Heathcare). HTB2-GFP–expressing cells were imaged at room temperature using a Nikon Ti2 microscope equipped with a Photometrics camera and 100X oil immersion objective lens. During time-lapse imaging, five z steps (1.2 µm) were acquired every 10 min. The exposure times used for bright-field and Htb2-GFP were 30msec and 10msec, respectively, with neutral-density filters transmitting 5% of light intensity. Images were acquired and data were analyzed using NIS Elements Viewer Version 4.50 software. Fiji software (NIH) was used to create final images with adjustment of brightness and contrast. All images were processed using ImageJ (National Institutes of Health). Z stacks were combined into a single maximum intensity projection to obtain the final images presented. Brightness and contrast were only adjusted on entire images.

### Whole genome sequencing and sequence analysis

Library preparation for whole genome sequencing was performed with the following protocol, adapted from Baym et al [2]. Cells from frozen glycerol stocks were inoculated into 100µL of rich medium, containing of 2% yeast extract, 1% peptone and 2% dextrose (YPD), in 96-well plates. Plates containing YPD cultures were incubated at 30°C for ∼24 hours or until all wells reached saturation. Cells were spheroplasted by mixing 5µL of YPD culture with 5µL of 5mg/mL 20T Zymolyaze (Sunrise Science) in a sorbitol/sodium phosphate buffer (1M Sorbitol, 100mM Sodium Phosphate pH 7.4, 20mM DL-Dithiothreitol (DTT)), and incubating for 30 minutes at 37°C. Tagmentation was performed directly on spheroplasts by mixing 1µL of spheroplasted cells with 0.25µL of TDE1 and 1.25µL of TD buffer (enzyme and buffer from Illumina) and incubating at 55°C for 15 minutes. The tagmented genome library was amplified with the addition of adapters by PCR - 2.5µL of tagmented DNA were mixed with 4.1µL sterile water, 11µL KAPA HotStart ReadyMix, and 2.2µL of each adapter primer. Fragment size selection was performed using Aline magnetic beads to select for fragments that are 300-500 residues long. The presence and size of genomic fragments was assayed by Qubit and Tapestation for quality control. Sequencing was performed on an Illumina NovaSeq with 150 base paired–end reads. Whole genome sequencing data were processed as described [16]. The Burrow-Wheeler Aligner (bio-bwa.sourceforge.net) was used to map DNA sequences to the S. cerevisiae reference genome r64, downloaded from Saccharomyces Genome Database (www.yeastgenome.org). The resulting SAM (Sequence Alignment/Map) file was converted to a BAM file, an indexed pileup format file, using the samtools software package (samtools.sourceforge.net). GATK (gatk.broadinstitute.org/hc/en-us) was used to realign local indels, and Varscan (varscan.sourceforge.net) was used to call variants. Mutations were identified using an in-house pipeline (github.com/koschwanez/mutantanalysis) written in Python. Mutations listed in figure 2C were present in 100% of reads in the given population.

## Supplementary figures

**Figure S1.**
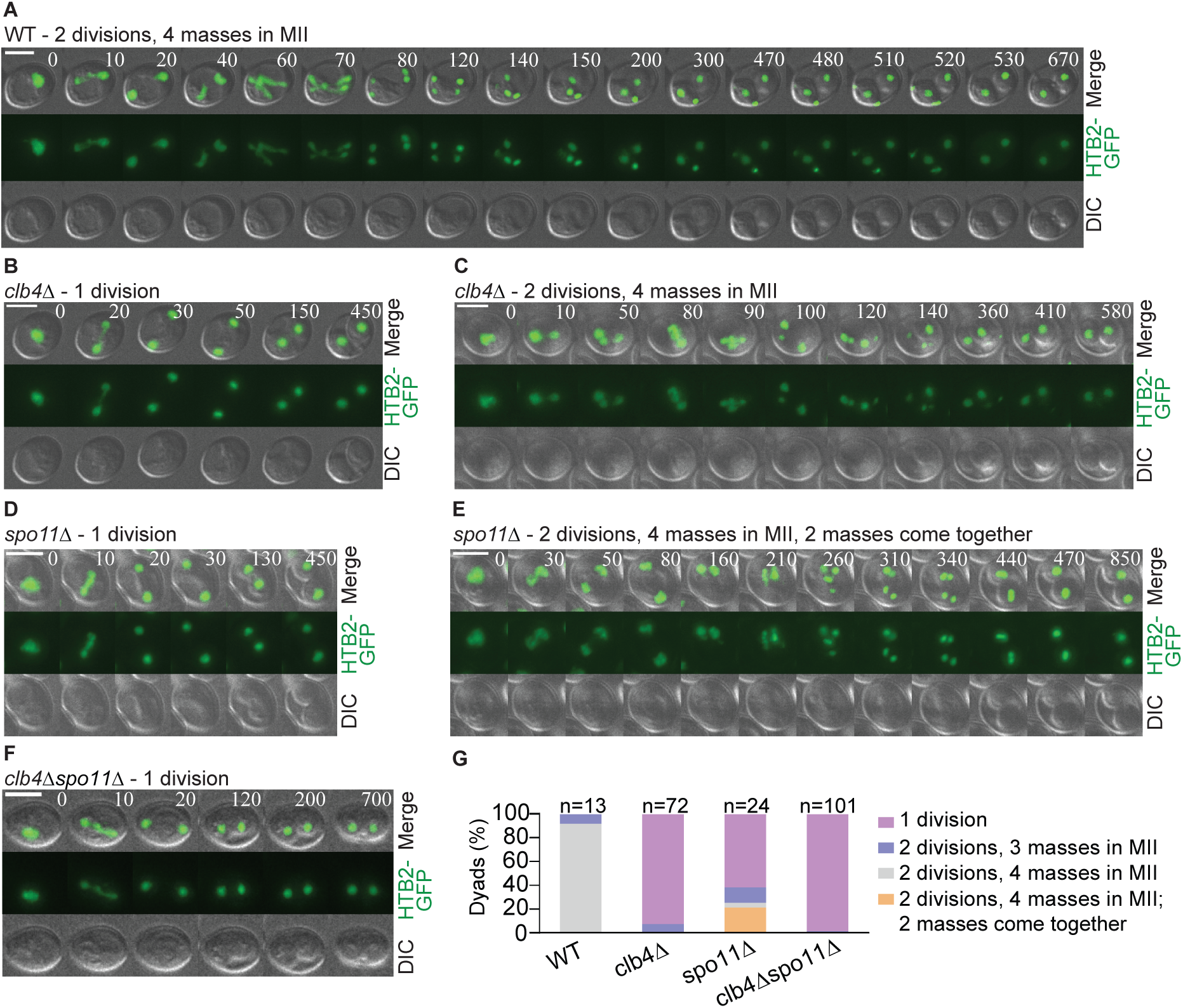
Meiotic chromatin mass divisions in *CLB4* and *SPO11* mutants. (A-F) Representative images from time lapse movies of sporulation with fluorescently labeled histone H2B (HTB2-GFP) in wild-type (WT, LY9981), *clb4*Δ (LY9982), *spo11*Δ (LY10028) and *clb4*Δ *spo11*Δ (LY9983). All deletions were homozygous. **(G)** Fraction of dyads observed forming in time lapse movies that underwent the listed division pattern. The number of dyads formed after 1 or 2 chromatin mass divisions was compared between the following strains using a chi-square test: wild-type (WT) vs. *clb4*Δ, *χ*^2^(*df* = 1*, N* = 85) = 57.13*, p <* 0.00001; *spo11*Δ vs *clb4*Δ *spo11*Δ, *χ*^2^(*df* = 1*, N* = 125) = 35.12*, p <* 0.00001.

**Figure S2.**
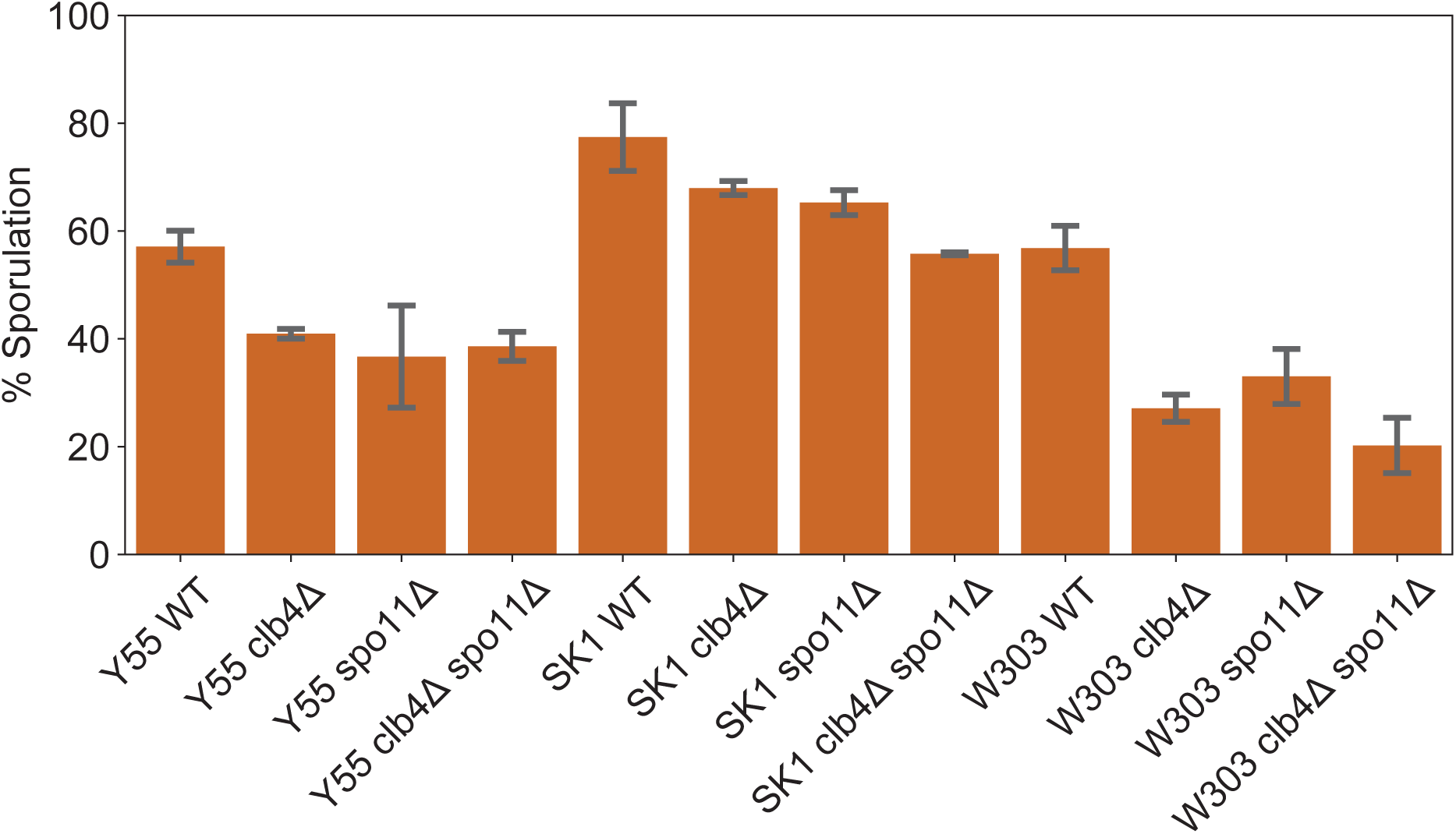
Sporulation fraction of *CLB4* mutants. The fraction of sporulated cells (asci) in the cultures assayed for figure 3.

**Figure S3.**
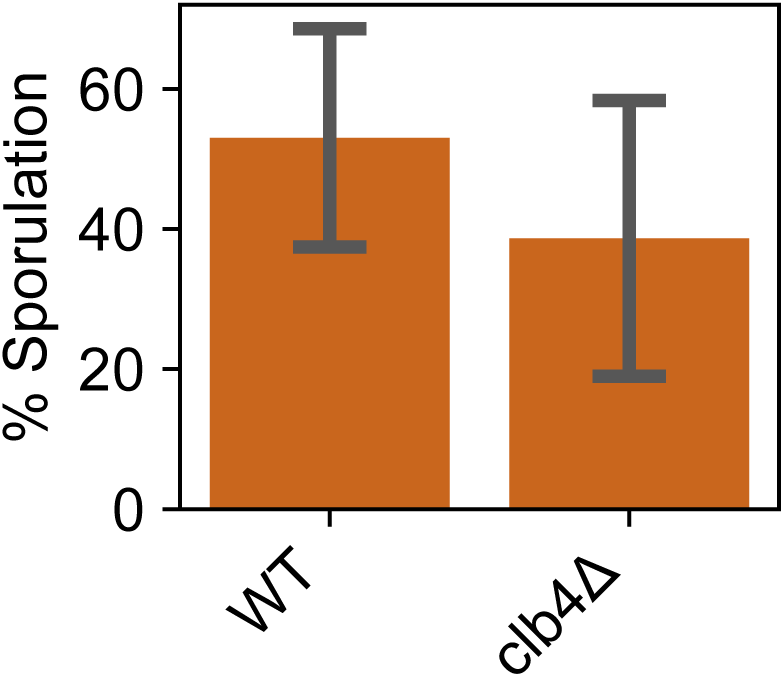
Sporulation fraction of meiotic time course cultures. The fraction of sporulated cells (asci) in the cultures assayed for figure 5.

**Figure S4.**
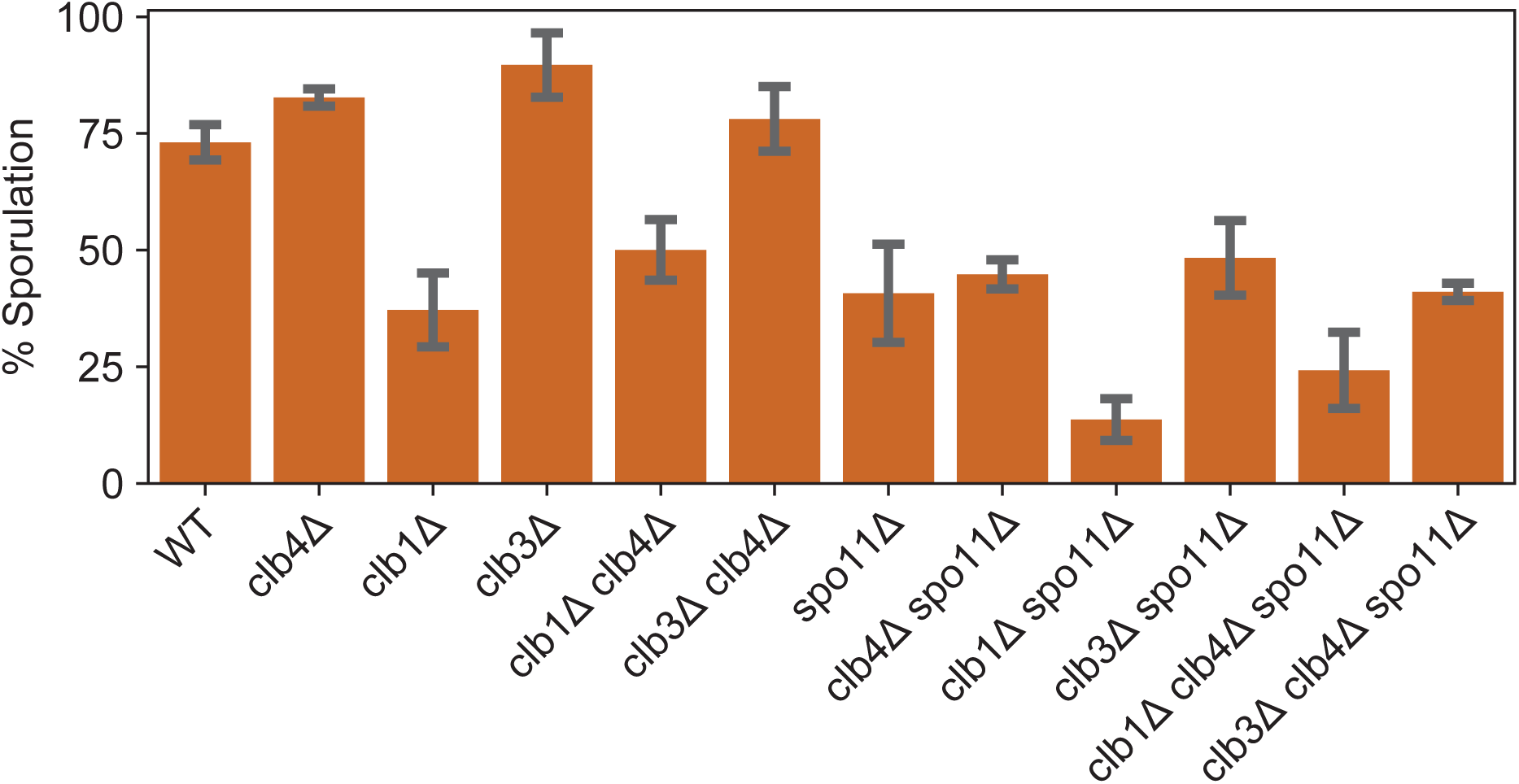
Sporulation fraction of *CLB* mutants. The fraction of sporulated cells (asci) in the cultures assayed for figure 7.

**Figure S5.**
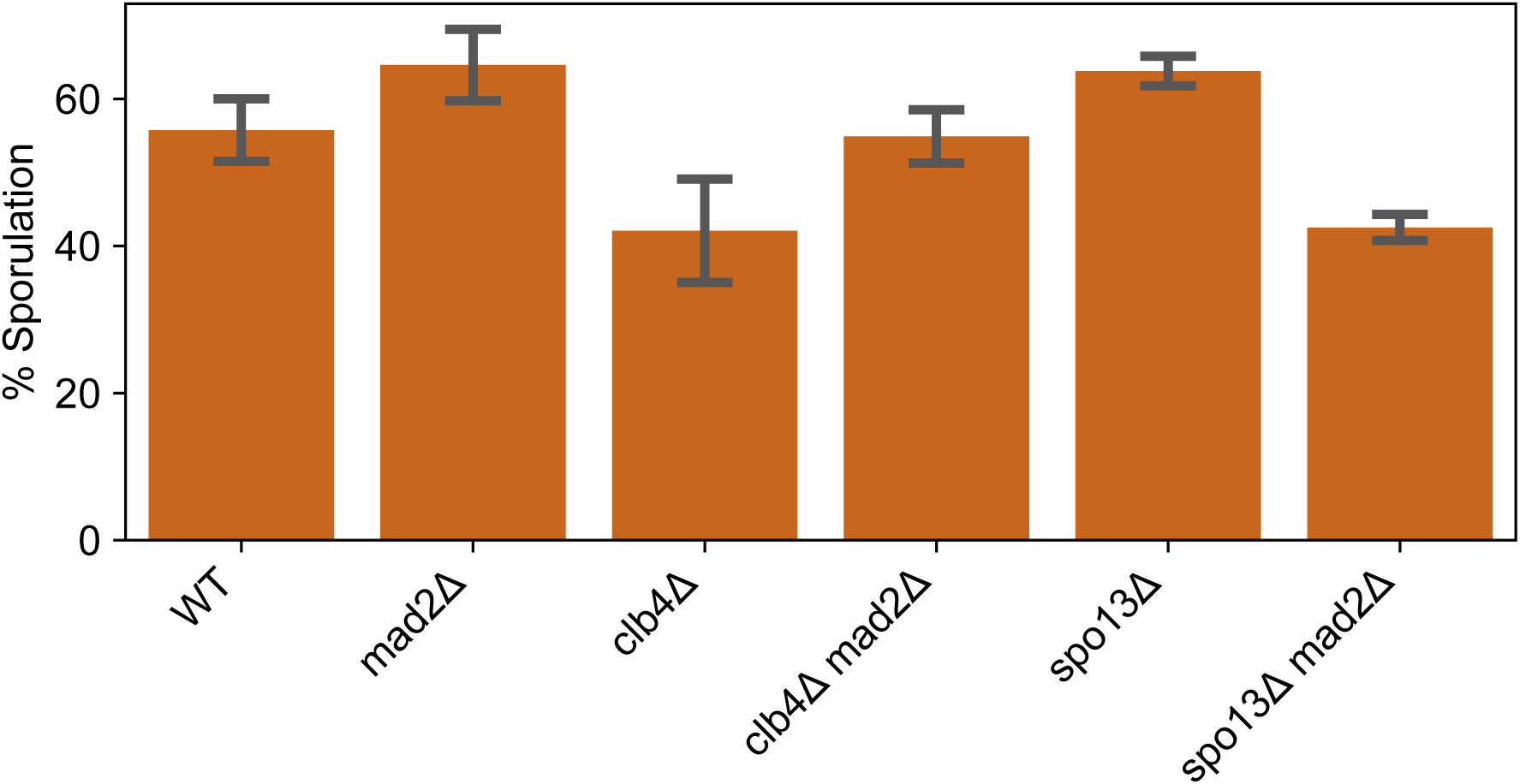
Sporulation fraction of *MAD2* mutants. The fraction of sporulated cells (asci) in the cultures assayed for figure 6.

